# Knockout of *cryptochrome 1* disrupts circadian rhythm and photoperiodic diapause induction in the silkworm, *Bombyx mori*

**DOI:** 10.1101/2024.05.13.593801

**Authors:** Hisashi Tobita, Takashi Kiuchi

## Abstract

Most insects enter diapause, a state of physiological dormancy crucial for enduring harsh seasons, with photoperiod serving as the primary cue for its induction, ensuring proper seasonal timing of the process. Although the involvement of the circadian clock in the photoperiodic time measurement has been demonstrated through knockdown or knockout of clock genes, the precise molecular mechanisms in this context remain unclear. In bivoltine strains of the silkworm, *Bombyx mori*, embryonic diapause is maternally controlled and affected by environmental conditions experienced by mother moths during embryonic and larval stages. Previous research highlighted the role of core clock genes, including *period* (*per*), *timeless* (*tim*), *Clock* (*Clk*) and *cycle* (*cyc*), in photoperiodic diapause induction in *B. mori*. In this study, we focused on another clock gene, *cryptochrome 1* (*cry1*), which functions as a photoreceptor implicated in photoentrainment of the circadian clock across various insect species. Phylogenetic analysis and conserved domain identification confirmed the presence of both *Drosophila*-type *cry* (*cry1*) and mammalian-type *cry* (*cry2*) genes in the *B. mori* genome, akin to other lepidopterans. Temporal expression analysis revealed higher *cry1* gene expression during the photophase and lower expression during the scotophase, with knockouts of core clock genes (*per*, *tim*, *Clk* and *cyc*) disrupting this temporal expression pattern. Using CRISPR/Cas9-mediated genome editing, we established a *cry1* knockout strain in p50T, a bivoltine strain exhibiting clear photoperiodism during both embryonic and larval stages. Although the wild-type strain displayed circadian rhythm in eclosion under continuous darkness, the *cry1* knockout strain exhibited arrhythmic eclosion, implicating *B. mori cry1* in the circadian clock feedback loop governing behavior rhythms. Females of the *cry1* knockout strain failed to induce photoperiodic diapause during both embryonic and larval stages, mirroring the diapause phenotype of the wild-type individuals reared under constant darkness, indicating that *B. mori* CRY1 contributes to photoperiodic time measurement as a photoreceptor. Furthermore, photoperiodic diapause induction during the larval stage was abolished in a *cry1*/*tim* double-knockout strain, suggesting that photic information received by CRY1 is relayed to the circadian clock. Overall, this study represents the first evidence of *cry1* involvement in insect photoperiodism, specifically in diapause induction.

**Highlights:**

- Knockouts of core clock genes disrupted the rhythmic expression of *cryptochrome 1* (*cry1*).
- A *cry1* knockout strain was established using CRISPR/Cas9.
- The *cry1* knockout strain lost its eclosion rhythm.
- Knockout of *cry1* disrupted photoperiodic diapause induction.
- Females of a *cry1*/*tim* double knockout strain produced only non-diapause eggs regardless of larval photoperiod.

**Graphical abstract:** 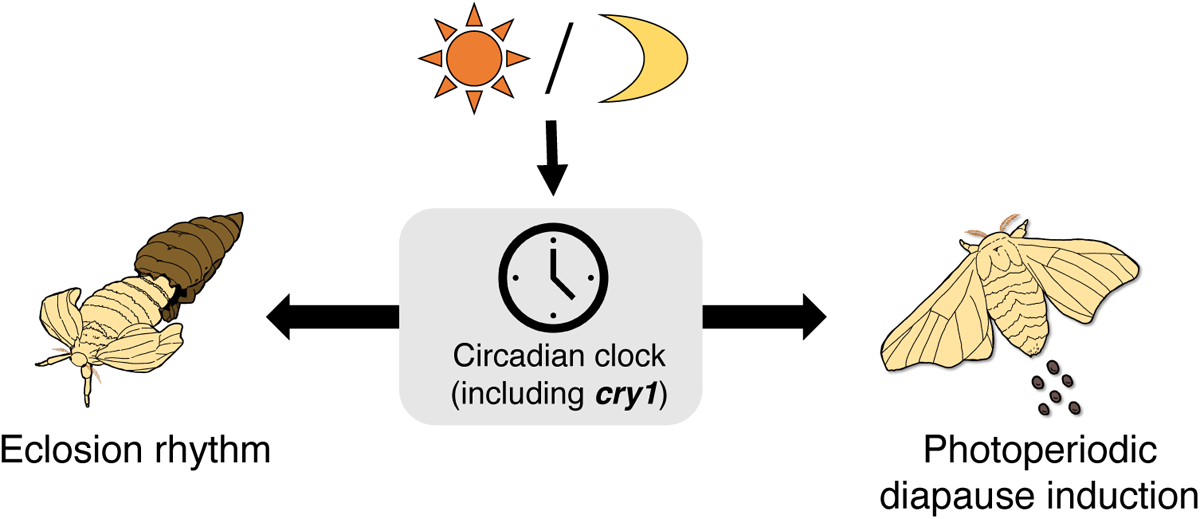

## 1. Introduction

Organisms exhibit endogenous circadian rhythms, approximately 24 h in duration, to anticipate and adapt to day–night cycles. The circadian clock is molecular basis generating the circadian rhythm and comprises autoregulatory transcriptional feedback loops. The clock’s regulatory mechanisms are well understood in the fruit fly, *Drosophila melanogaster* (Beer and Helfrich-Förster, 2020), where a core feedback loop, primarily driven by four clock genes, governs the central circadian clock. CLOCK (CLK) and CYCLE (CYC) form a heterodimer and bind to E-box motifs to activate *period* (*per*) and *timeless* (*tim*) transcription (Allada et al., 1998; Rutila et al., 1998). The encoded PER and TIM proteins dimerize and enter the nucleus to suppress CLK and CYC heterodimer transcriptional activity (Darlington et al., 1998). Therefore, CLK and CYC are designated positive elements, whereas PER and TIM are considered negative elements. Additionally, rhythmic expression of *Clk* is positively regulated by *par domain protein 1* (*pdp1*) and negatively regulated by *vrille* (*vri*), further reinforcing clockwork oscillation (Cyran et al., 2003; Glossop et al., 2003).

The circadian clock is entrained with environmental signals, such as photoperiod. *Drosophila cryptochrome* (*cry*) gene plays a role in photic entrainment of the circadian clock (Emery et al., 1998, 2000; Stanewsky et al., 1998). *Drosophila* cryptochrome protein (CRY) is a blue-light photoreceptor that directly interacts with TIM protein and promote its degradation in a light-dependent manner (Ceriani et al., 1999). In insects, two types of *cry* genes exist: *Drosophila*-type *cry* (*cry1* or *cry-d*) and *mammalian*-type *cry* (*cry2* or *cry-m*). Evolutionary gene duplication and loss have resulted in *cry* gene diversification among insect species. For example, *D*. *melanogaster* possesses only *cry1*, whereas the western honeybee, *Apis melifera*, the fire ant, *Solenopsis invicta*, and the red flour beetle, *Tribolium castaneum* possess only *cry2* (Rubin et al., 2006; Zhu et al., 2006; Ingram et al., 2012). Conversely, some insects possess both *cry1* and *cry2*, e.g., the mosquito species *Aedes aegypti* and *Anopheles gambiae*, the pea aphid, *Acyrthosiphon pisum*, and the monarch butterfly, *Danaus plexippus* (Zhu et al., 2006; Yuan et al., 2007; Cortés et al., 2010). In insects, CRY1 serves as a blue-light photoreceptor involved in circadian clock photoentrainment (Emery et al., 1998, 2000; Stanewsky et al., 1998). In contrast, CRY2 lacks photosensitivity, exhibiting a clockwork function as a transcriptional repressor that inhibits CLK/CYC activity along with other negative elements, such as PER and TIM. *In vitro* experiments using *Drosophila* Schnider2 (S2) cells have shown that CRY2 from various insect species, including *T*. *castaneum, A*. *gambiae, A*. *melifera,* the Chinese oak silk moth *Antheraea pernyi,* and the two-spotted cricket, *Gryllus bimaculatus* can inhibit CLK/CYC-mediated transcription (Zhu et al., 2006; Yuan et al., 2007; Tokuoka et al., 2017). CRY2 from *D. plexippus*, has been shown to function as a repressor in DpN1 cells derived from *D. plexippus* embryos (Zhu et al., 2008). These results indicate that CRY2 is involved in the negative feedback loop with PER and TIM in non-drosophilid insects.

Most insects survive winter by entering diapause, a physiologically dormant state triggered by environmental cues. Photoperiod, the ratio of day length to night length, serves as the most reliable signal for anticipating seasonal change. Insects measure photoperiod and enter diapause at correct timing. The involvement of the circadian clock in this photoperiodic time measurement system has been indicated in several studies involving gene manipulation techniques, such as RNAi and genome editing, applied to clock genes affecting photoperiodic diapause induction in numerous insect species (Ikeno et al., 2010, 2011, 2013; Meuti et al., 2015; Mukai and Goto, 2016; Kotwica-Rolinska et al., 2017, 2022; Iiams et al., 2019; Tamai et al., 2019; Goto and Nagata, 2022). These studies have highlighted the circadian clock’s role in insect photoperiodic diapause induction, although the molecular mechanisms remain poorly understood. The involvement of *cry2* in photoperiodic diapause induction has been reported in various insect orders, including Hemiptera (e.g., the linden bug *Pyrrhocoris apterus* and the bean bug *Riptortus pedestris*), Diptera (e.g., the northern house mosquito, *Culex pipiens*) and Lepidoptera (e.g., *D. plexippus*), whereas the role of *cry1* in photoperiodism remains unclear. CRY1 has been believed to be involved in the photoperiodic time measurement system as a photoreceptor (Saunders, 2012). In the cricket species *Modicogryllus siamensis*, RNAi targeting *cry1* affects the photoperiodic response in nymphal development (Ueda et al., 2018); however cricket *cry1* functions as a negative element, not as a photoreceptor (Tokuoka et al., 2017). Therefore, no study has definitively demonstrated the involvement of photosensitive *cry1* in insect photoperiodism.

The silkworm, *Bombyx mori* exhibits clear photoperiodism in the diapause induction. In bivoltine strains, diapause in progeny is determined by temperature and photoperiodic conditions experienced by mother moths during embryonic and larval stages (Kogure, 1933; Egi et al., 2014). Diapause hormone (DH) synthesis and release play crucial roles in diapause induction. DH, synthesized in the suboesophageal ganglion and released from the corpus cardiacum into the hemolymph during pupal to adult development in diapause egg producers, acts on DH receptors in developing ovaries, inducing diapause (Takeda and Ogura, 1976; Sato et al., 1998). Conversely, in non-diapause egg producers, DH release is suppressed through GABAergic and corazonin signaling pathways (Shimizu et al., 1989; Tsuchiya et al., 2020). Additionally, temperature-dependent transcriptional change of the plasma membrane GABA transporter gene in pupal brain‒ suboesophageal ganglion complex is implicated in embryonic temperature-dependent diapause induction (Tsuchiya et al., 2020). Similar to the aforementioned studies on other insects, the involvement of circadian clock genes in the photoperiodic diapause induction has been reported in *B*. *mori* (Ikeda et al., 2021; Tobita and Kiuchi, 2022). For instance, Ikeda et al. (2021) found that *per* knockout in the bivoltine strain Kosetsu resulted in non-diapause egg production in most knockout females, even with rearing under short photoperiod (diapause condition) during larval stage. Previously, we established knockout strains of *per*, *tim*, *Clk*, and *cyc* in the bivoltine strain p50T, observing loss of responsiveness to photoperiod during embryonic and larval stages in knockout females, resulting in non-diapause egg production regardless of photoperiodic conditions (Tobita and Kiuchi, 2022). Furthermore, Homma et al., (2022), having knocked out not only *per*, *tim*, *Clk* and *cyc* but also *cry1* and *cry2*, reported effects on temperature-dependent diapause induction, finding disruption in *cry2* but not *cry1* knockout. In recent years, studies using the *cry1* knockout strain have revealed *cry1*’s role in hatching rhythm, immunity, cell growth, and hormonal control of development (Qiu et al., 2022, 2023a, 2023b; Yuan et al., 2023). However, the involvement of *cry* genes in *B. mori* photoperiodism remains unclear. To elucidate the molecular mechanisms of the circadian clock in photoperiodic time measurement for diapause induction of *B. mori*, we focused on *cry1*, a constituent of the circadian clock’s canonical feedback loop. Specifically, we employed CRISPR/Cas9-mediated knockout of the *cry1* gene to investigate its effect on circadian rhythm and photoperiodic diapause induction.

## 2. Materials and methods

### 2.1. Silkworm strains

The inbred bivoltine strain p50T, derived from the Daizo strain, is maintained in our laboratory, and was used in this study. Larvae were fed with mulberry leaves or artificial diet Silkmate PS (NOSAN) and reared at 25℃ under a daily 12:12 h light:dark cycle (12L12D). To obtain non-diapause eggs for microinjection, eggs were maintained at 18℃ and larvae were subsequently reared under long-day conditions (15L9D or 20L4D).

### 2.2. Phylogenetic analysis

Amino acid sequences of CRY from vertebrates and insects were retrieved from GenBank. These sequences were aligned using ClustalW, and a phylogenetic tree was constructed employing maximum likelihood methods based on the Jones‒Taylor‒Thornton matrix model using MEGA X software (Kumar et al., 2018). Bootstrap tests were performed with 1000 replicates. Accession numbers of protein sequences used in the phylogenetic analysis are listed in Table S1.

### 2.3. Quantitative real-time PCR

Non-diapause eggs were maintained at 25℃ under continuous darkness (DD) until hatching, followed by rearing under 12L12D with artificial diet during the larval stage. Larval sampling was performed on the third day of the fifth instar. Larvae were immediately frozen at zeitgeber time (ZT) 1, ZT5, ZT9, ZT13, ZT17, and ZT21 under 12L12D conditions and stored at −80℃. Each larvae’s whole head was homogenized in TRIzol reagent (Invitrogen), and total RNA was extracted and subsequently used to synthesize cDNA using the TaKaRa RNA PCR^TM^ Kit (AMV) Ver.3.0 (TaKaRa). Quantitative real-time PCR (qPCR) was performed using the KAPA^TM^ SYBR FAST qPCR Kit (KAPA Biosystems) and StepOnePlus^TM^ Real-Time PCR System (Applied Biosystems). Transcript levels were determined using the 2^-ΔΔCt^ method. The genes *actinA3* and *A4* served as references for normalizing the expression levels of the *cry1* gene. Primers used for qPCR are listed in Table S2. To assess the statistical significance of temporal differences in *cry1* gene expression, one-way ANOVA and the Tukey-Kramer *post hoc* test were employed. Two-way ANOVA and Bonferroni’s *post hoc* test were used to evaluate significant differences in *cry1* gene expression between the wild-type and each knockout strain.

### 2.4. CRISPR/Cas9-mediated gene knockouts

A unique single guide RNA (sgRNA) was designed for the target gene, *cry1* (KWMTBOMO10548), using CRISPRdirect (Naito et al., 2015: https://crispr.dbcls.jp/) based on their coding sequences. The sgRNA was synthesized *in vitro* following the method of Bassett et al., (2014) using the primers listed in Table S2. Cas9 protein (600 ng/µL final concentration; Nippon gene) and sgRNA (150 ng/µL final concentration) were mixed in injection buffer [100 mM KOAc, 2 mM Mg (OAc)_2_, 30 mM HEPES-KOH; pH 7.4] and injected into non-diapause eggs within 2–4 h postoviposition.

### 2.5. Establishment of knockout strains

Hatched G_0_ (injected generation) individuals were raised to adulthood and subsequently crossed with wild-type moths to obtain G_1_ eggs. Genomic DNA was extracted from a leg of each G_1_ individual using the HotSHOT method (Truett et al., 2000). Genomic PCR was performed using KOD One (TOYOBO) with the primers listed in Table S2. The PCR reaction proceeded as follows: 35–40 cycles of 98℃ for 10 s, 60℃ for 5 s, and 68℃ for 5 s/kb. Mutations were identified through the heteroduplex mobility assay (HMA) using the MultiNA (SHIMADZU) microchip electrophoresis system (Ota et al., 2013; Ansai et al., 2014). G_1_ moths exhibiting identical mutations, confirmed via band patterns in HMA, were crossed to generate a homozygous knockout strain. Genomic PCR products were sequenced using the ABI3130xl genetic analyzer (Thermo Fisher Scientific) to determine mutation patterns. A *cry1*/*tim* double-knockout strain was established through crossing between *cry1^Δ8^*and *tim^In11^* strains (Tobita and Kiuchi, 2022). Homozygous knockouts were selected through the HMA and subsequently crossed with each other from the F_2_ generation onward.

### 2.6. Recording and analysis of eclosion rhythm

Eclosion rhythm was recorded under DD at 25℃, with insects reared under 12L12D conditions until recording. Four days after spinning (2days after pupation), pupae were removed from cocoons and placed in a box. They were transferred to constant darkness at lights-off (ZT12) nine days after spinning and recording was started. Eclosion was recorded at 20 min intervals using a compact digital camera (Tough TG-6, Olympus). A dim red LED light (Kaito denshi; 660 nm) served as the light source for recording, maintaining the light intensity of approximately 1.0 mW/m^2^. The degree of rhythmicity in eclosion was analyzed using the parameter R (Winfree, 1970), with R values calculated as follows: first, the gate period was determined as the 8 h period with the highest number of eclosions; second, the number of insects eclosed outside the gate period was divided by the number of insects eclosed during the gate period and multiplied by 100. R ≤ 60 indicated statistically rhythmic eclosion, 60 < R ≤ 90 represented weakly rhythmic eclosion, and R > 90 showed arrhythmic eclosion.

### 2.7. Diapause phenotyping

Non-diapause eggs were used, and hatched larvae were fed with artificial diet Silkmate PS to standardize nutritional conditions. To observe the effect of the gene knockout on photoperiodic diapause induction during the larval stage, eggs were maintained at 25℃ under DD, and hatched larvae were reared under long-day (20L4D) or short-day (8L16D) conditions. To observe the effect during the embryonic stage, eggs were maintained at 18℃ under DD or continuous light (LL), and hatched larvae were subsequently reared under 12L12D conditions. Female moths resulting from these conditions were crossed with male moths, and the oviposited eggs were maintained at 25℃ for 7–8 days. The diapause phenotype of each egg was assessed as previously described (Tobita and Kiuchi, 2022). Ommochrome-pigmented black eggs were classified as diapause eggs, whereas unpigmented and developed eggs until the head pigmentation stage were categorized as non-diapause eggs. Uncolored and undeveloped eggs were considered unfertilized and were excluded from further calculations. The total number of diapause and non-diapause eggs in each batch, excluding unfertilized eggs, was counted. Batches with > 90% diapause or non-diapause eggs were classified as diapause or non-diapause diapause or non-diapause batches, respectively, whereas batches with ≥10% mixed eggs were categorized as mix batches. Subsequently, the diapause phenotype of each strain was evaluated based on the ratio of diapause, non-diapause and mix batches to the total number of batches.

## 3. Results

### 3.1. CRY proteins in B. mori

*Bombyx mori cry1* and *cry2* were previously identified from a new genome assembly by Kawamoto et al., (2019). Both amino acid sequences, based on the predicted gene models, commonly possess the DNA-photolyase domain and the flavin adenine dinucleotide (FAD) binding domain, characteristic motifs in CRYs (Fig.1A). RD-2a, RD-1, RD-2b, and inhibition of CLOCK-ARNTL-mediated transcription (ICAT) domains necessary for repressing CLK/CYC-mediated transcription are highly conserved in *B. mori* CRY2 but not CRY1 (Fig. 1A; Hirayama and Sassone-Corsi, 2005). Additionally, we identified a nuclear localization signal (NLS) in the RD-2b domain of *B. mori* CRY2 (Fig. S1). Phylogenetic analysis of CRY proteins in vertebrates and insects revealed two distinct clusters (Fig. 1B): one consisting of *Drosophila*-type CRYs (CRY1 or CRY-d) and the other of mammalian-type CRYs (CRY2 or CRY-m). *Bombyx mori* CRY1 and CRY2 belong to the *Drosophila*-type and mammalian-type CRY groups, respectively. *Bombyx mori* CRY1 formed a clade with other lepidopteran CRY1s, such as *D. plexippus* and *A. pernyi* CRY1s, whereas *B. mori* CRY2 formed a clade with other lepidopteran CRY2s. Based on these results, we conclude that *B. mori cry1* and *cry2* are orthologous genes of *cry-d* and *cry-m*, respectively.

**Figure 1.**
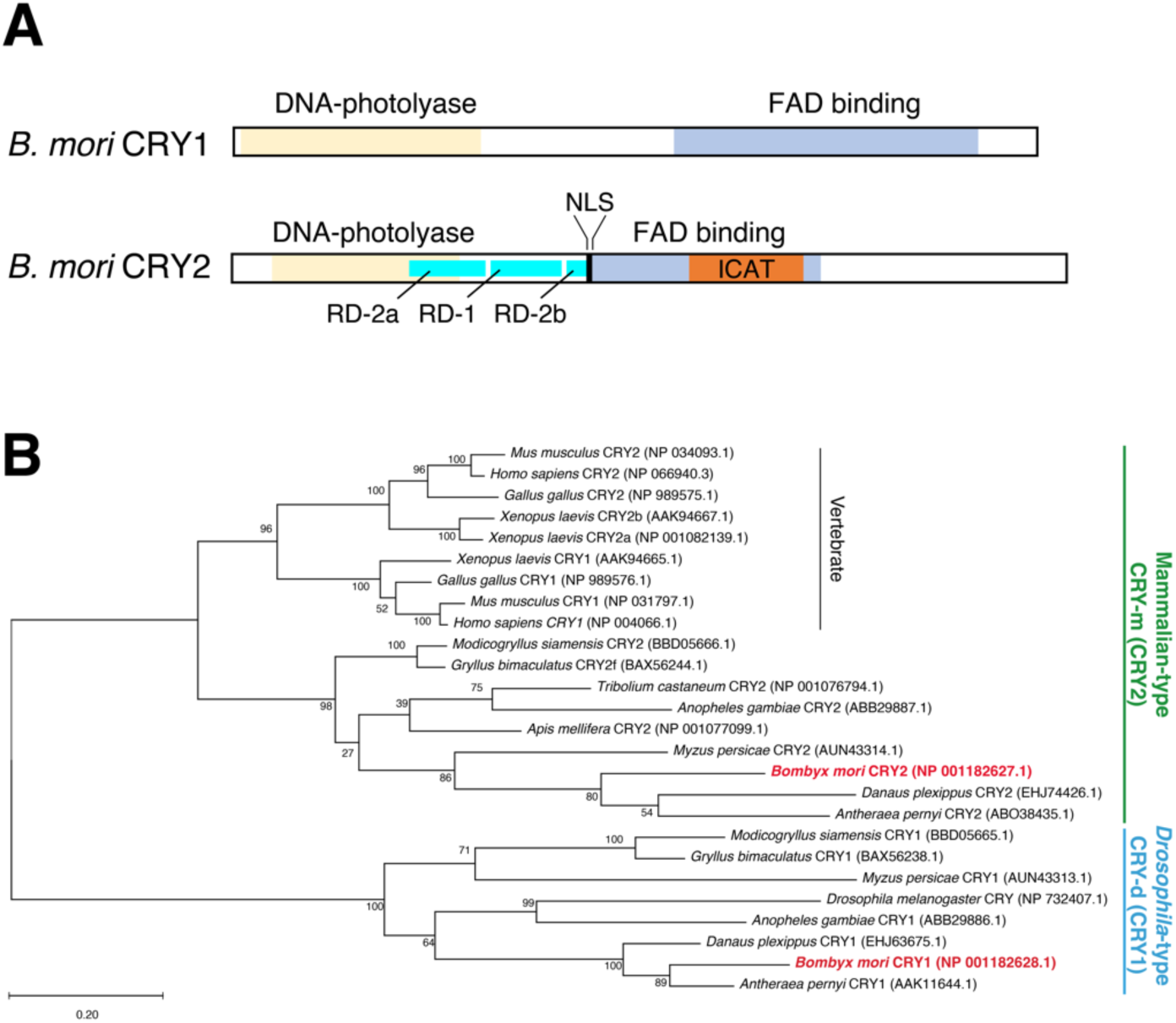
Predicted protein structures of *B. mori* CRY proteins and phylogenetic analysis. (A) Domain structures of *B. mori* CRY1 and CRY2. Functional domains are represented by colored boxes: DNA-photolyase (cream), FAD binding (blue), RD (cyan), NLS (black), and ICAT (orange) domains. (B) Phylogenetic analysis of CRY proteins in various insects and mammals. CRY amino acid sequences were obtained from GenBank (accession numbers appear next to species names; Table S1). Numbers on nodes represent the bootstrap values with 1000 replicates.

### 3.2. Expression pattern of the cry1 gene

In the wild-type strain, *cry1* exhibited a clear temporal expression change, peaking during the early photophase (ZT1 and 5) and gradually decreasing thereafter (Fig. 2; *p* < 0.001, one-way ANOVA). The relative expression levels of *cry1* fluctuated approximately 1.8-fold within a day. As we had previously established knockout strains of the core clock genes *per*, *tim*, *Clk* and *cyc*, designated as *per^Δ2^*, *tim^In11^*, *Clk^Δ5^*, and *cyc^In2^*, respectively (Tobita and Kiuchi, 2022), we also measured mRNA levels of *cry* genes in these knockout strains (Fig. 2). In the knockout strain of the negative element *per^Δ2^*, *cry1* expression was decreased during the photophase, (*p* < 0.001, two-way ANOVA). In the *tim^In11^*strain, *cry1* expression was also decreased during the photophase (*p* < 0.01, two-way ANOVA). Knockout strains of the positive elements *Clk^Δ5^*and *cyc^In2^* exhibited significant upregulation of *cry1* expression at all time points (*p* < 0.001, two-way ANOVA).

**Figure 2.**
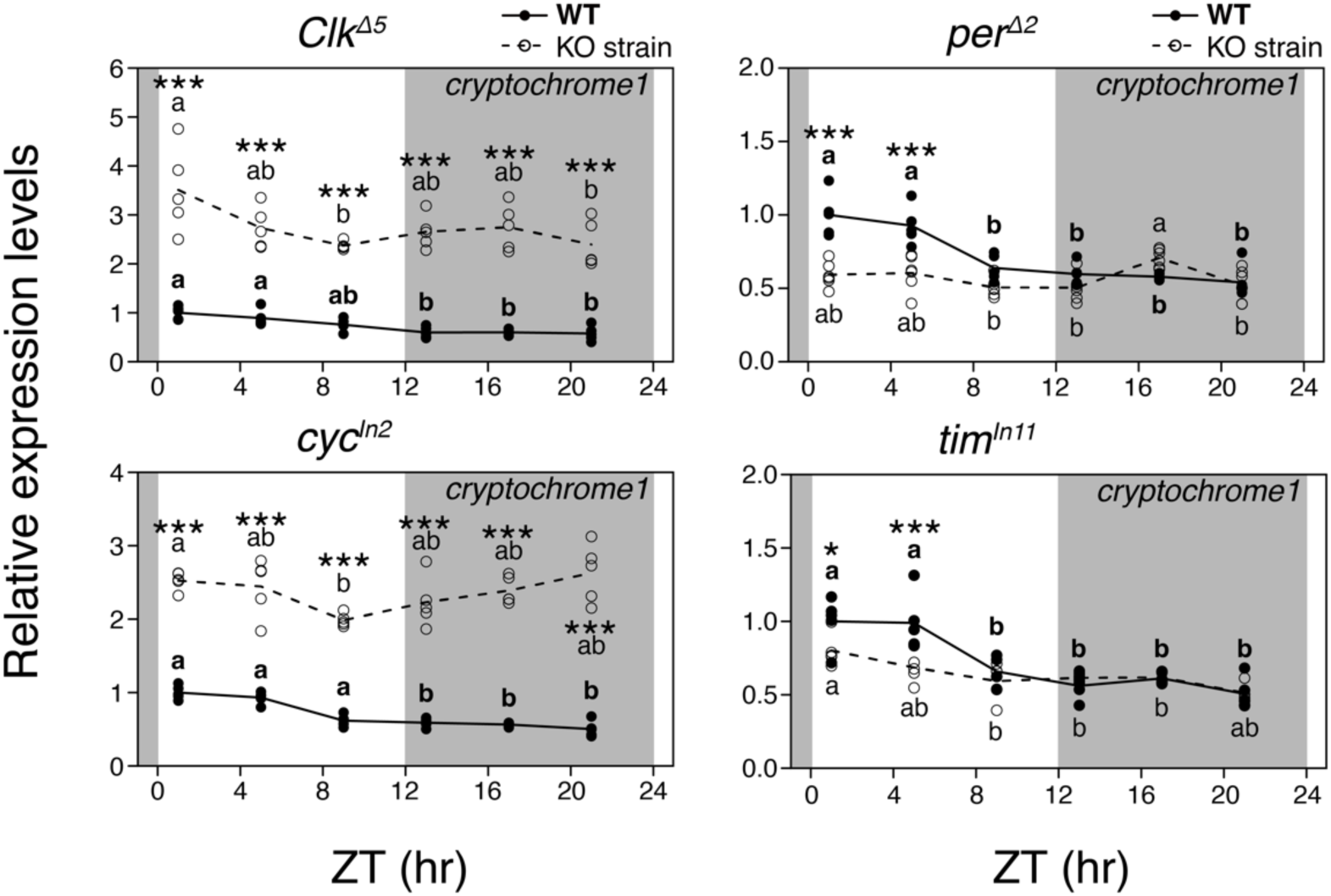
Temporal expression patterns of *cry1* in core clock gene knockout strains: *per^Δ2^*, *tim^In11^*, *Clk^Δ5^* and *cyc^In2^*. Expression levels were examined via qPCR using the heads of day-three fifth-instar larvae under 12L12D conditions. Relative expression levels (means of the wild-type at ZT1 = 1) of wild-type (closed circles) and *cry1^Δ8^* (open circles) strains are indicated by solid and broken lines, respectively (n = 5 or 6). The genes *actin A3* and *A4* genes served as references for normalization. Shaded areas indicate the scotophase. Lower case letters and asterisks indicate significant differences in relative expression levels throughout the day (one-way ANOVA and Tukey–Krammer *post hoc* tests) and between wild-type and *cry1^Δ8^* strains at each time point (two-way ANOVA and Bonferroni’s *post hoc* test), respectively (\**p* < 0.05, \*\**p* < 0.01, and \*\*\**p* < 0.001).

### 3.3. Establishment of a cry1 knockout strain

To investigate the involvement of the *cry1* gene in photoperiodic diapause induction, CRISPR/Cas9-mediated knockout was performed. The inbred strain p50T was chosen for the gene knockout as this strain responds to both embryonic and larval photoperiods (Egi et al., 2014; Tobita and Kiuchi, 2022). We designed sgRNA targeting the 4th exon of *cry1* and established a knockout strain, designated as *cry1^Δ8^* (Fig. 3). The *cry1^Δ8^* strain possessed an 8 bp deletion at the target site, potentially producing a truncated CRY1 protein lacking a part of the DNA-photolyase domain and the entire FAD binding domain (Fig. 3), suggesting that the *cry1^Δ8^* strain is a null mutant.

**Figure 3.**
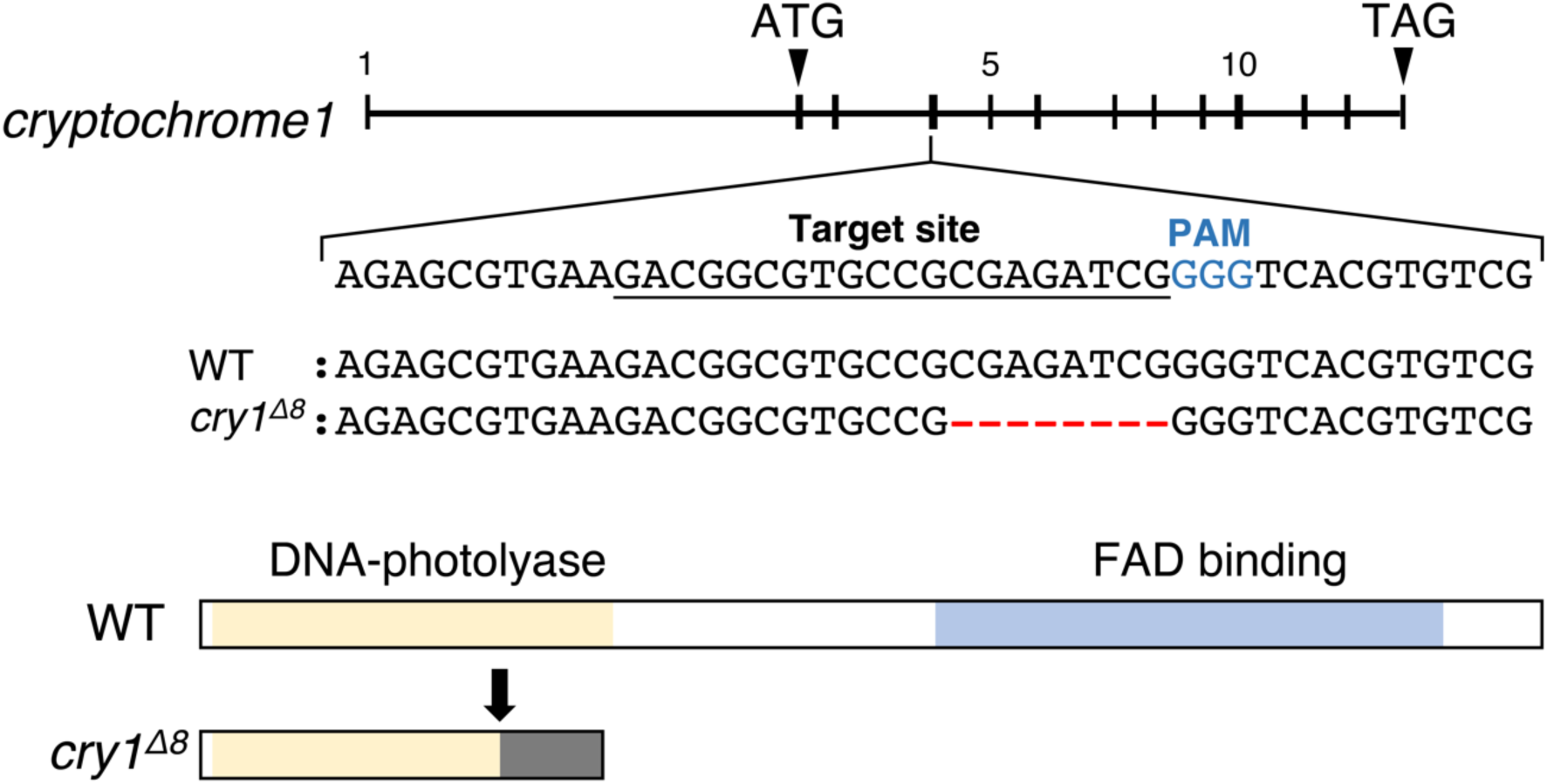
CRISPR/Cas9-mediated *cry1* gene knockout. Genomic structure of *cry1*, with the sgRNA target site (underlined) and the protospacer adjacent motif site (blue letters) shown at the top. Alignment of nucleotide sequences surrounding the sgRNA target from the wild-type and the *cry1* knockout strain (*cry1^Δ8^*) strains are shown at the center. Red letters indicate 8 bp deletions. Predicted protein structures are shown at the bottom. Black arrow indicates the mutation site. Functional domains are represented by colored boxes: DNA-photolyase (cream) and FAD binding (blue) domains.

### 3.4. Eclosion rhythm in the cry1 knockout strain

To assess the impact of *cry1* gene knockout on circadian rhythm, we investigated eclosion rhythm, as the timing of eclosion in *B. mori* is regulated by the circadian clock (Ikeda et al., 2019, 2021; Nartey et al., 2021). In the wild-type strain, moths emerged intensively from subjective late night to early day under constant darkness (Fig. 4A). The R value was 20.9, indicating rhythmic eclosion in the wild-type strain. Conversely, moths of the *cry1^Δ8^* strain eclosed sporadically throughout the recording period, with the calculated R value of 94.5, indicating arrhythmicity (Fig.4B). Therefore, we concluded that the *cry1* knockout strain had lost eclosion rhythm.

**Figure 4.**
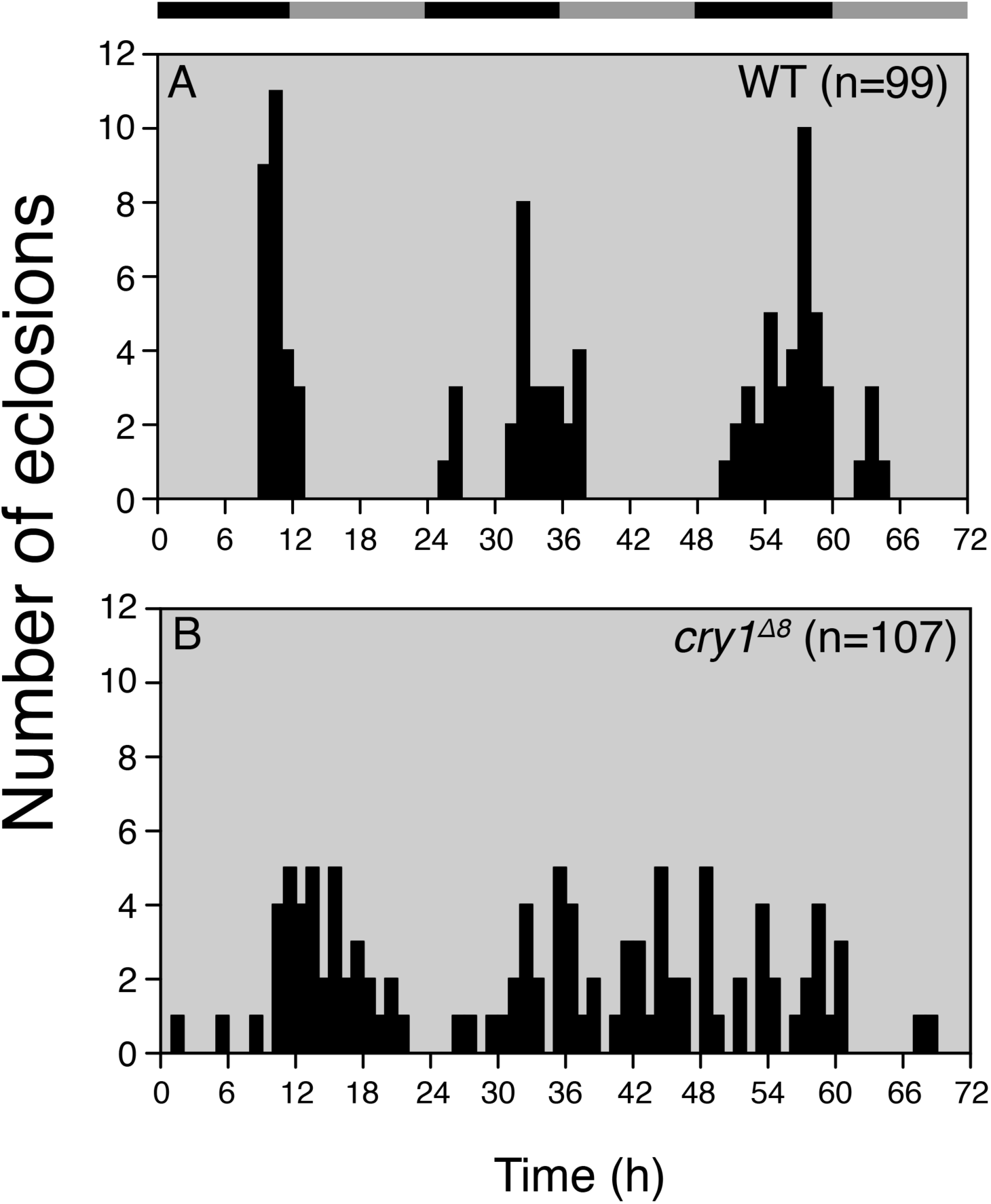
Distribution of adult eclosions in wild-type and *cry1^Δ8^* strains under continuous darkness at 25℃. Black and gray horizontal bars indicate subjective night and day, respectively.

### 3.5. Effect of cry1 gene knockout on photoperiodic diapause induction

To analyze the effect of *cry1* gene knockout on photoperiodic diapause induction during the larval stage, we incubated eggs at 25℃ under DD and then reared larvae under long-day (20L4D) or short-day (8L16D) conditions. As previously reported by Tobita and Kiuchi (2022), wild-type females exhibited clear photoperiodism, with all females producing diapause eggs under short-day conditions and almost all females producing non-diapause eggs under long-day conditions (Fig. 5). Upon *cry1* knockout, 80% of females produced diapause eggs, even when reared under long-day conditions. Subsequently, we examined the involvement of the *cry1* gene in photoperiodic diapause induction during the embryonic stage. Eggs were incubated at 18℃ under DD or LL until hatching, after which larvae were reared under 12L12D conditions. When wild-type females experienced DD during the embryonic stage, 55%, 35%, 10% produced diapause eggs, non-diapause eggs, mixed eggs, respectively (Fig. 6). Conversely, wild-type females experiencing LL during the embryonic stage produced solely diapause eggs (Fig. 6). However, in the *cry1^Δ8^* strain, all females produced non-diapause eggs regardless of embryonic photoperiod (Fig. 6). CRY1 is considered the circadian photoreceptor in insects; therefore, we investigated the diapause phenotype of the wild-type strain reared under constant darkness. Consequently, when embryos were maintained at 25℃, 90% of resultant females produced diapause eggs, whereas when they were maintained at 18℃, all resultant females produced non-diapause eggs (Fig. 7). To examine the hierarchical relationship between *cry1* and the circadian clock, a *cry1*/*tim* double-knockout strain was generated via crossing. We investigated its diapause phenotype during the larval stage: eggs were maintained at 25℃ under DD, and larvae were reared under short-day or long-day conditions. Although females of the *cry1* knockout strain predominantly produced diapause eggs regardless of larval photoperiod (Fig. 5), females of the *cry1*/*tim* double knockout strain produced solely non-diapause eggs (Fig. 8).

**Figure 5.**
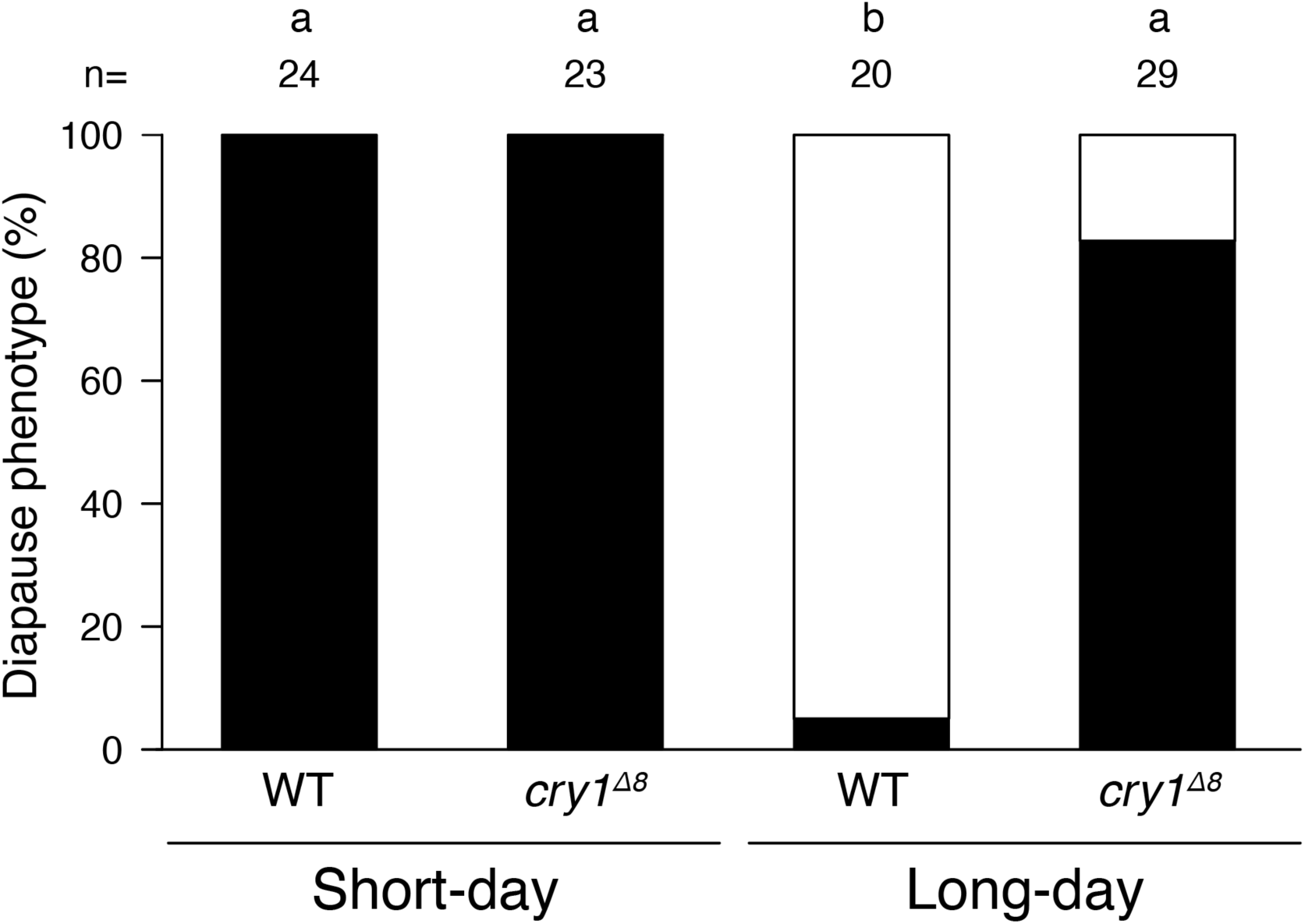
Effect of *cry1* gene knockout on photoperiodic diapause induction during the larval stage. Eggs were incubated under continuous darkness at 25 ℃, and hatched larvae were reared under short-day (8L16D) or long-day (20L4D) conditions. Black- and white-shaded parts show the proportions of females ovipositing diapause and non-diapause eggs, respectively. No female oviposited both diapause and non-diapause eggs in the same batch. Different letters above columns indicate statistically significant differences (Tukey-type multiple comparisons for proportions).

**Figure 6.**
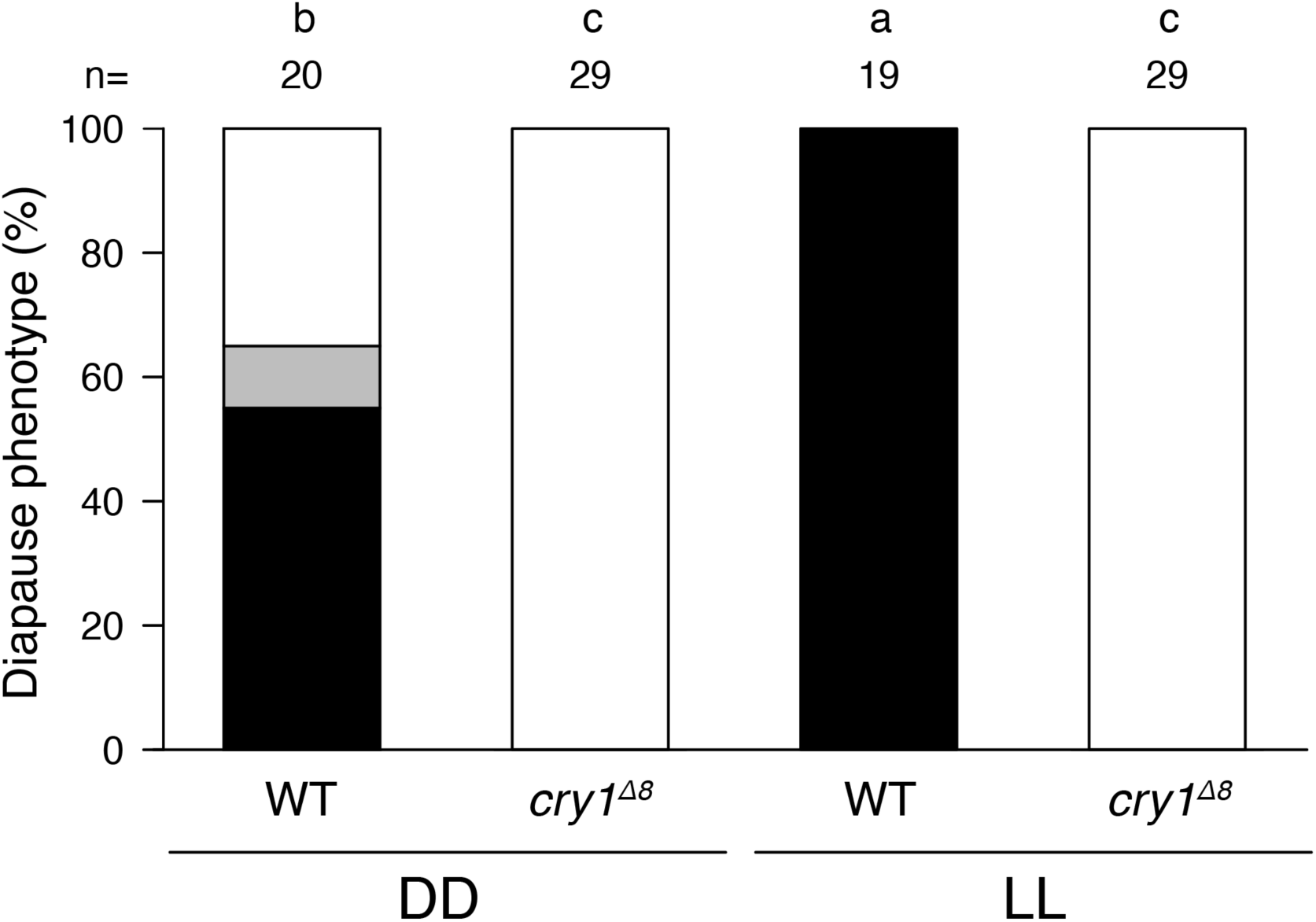
Effects of *cry1* gene knockout on photoperiodic diapause induction during the embryonic stage. Eggs were incubated under continuous darkness (DD) or continuous light (LL) conditions at 18℃ until hatching. Hatched larvae were reared under 12L12D. Black-, gray- and white-shaded parts show the proportions of females ovipositing diapause, mix and non-diapause eggs, respectively. Different letters above columns indicate statistically significant differences (Tukey-type multiple comparisons for proportions).

**Figure 7.**
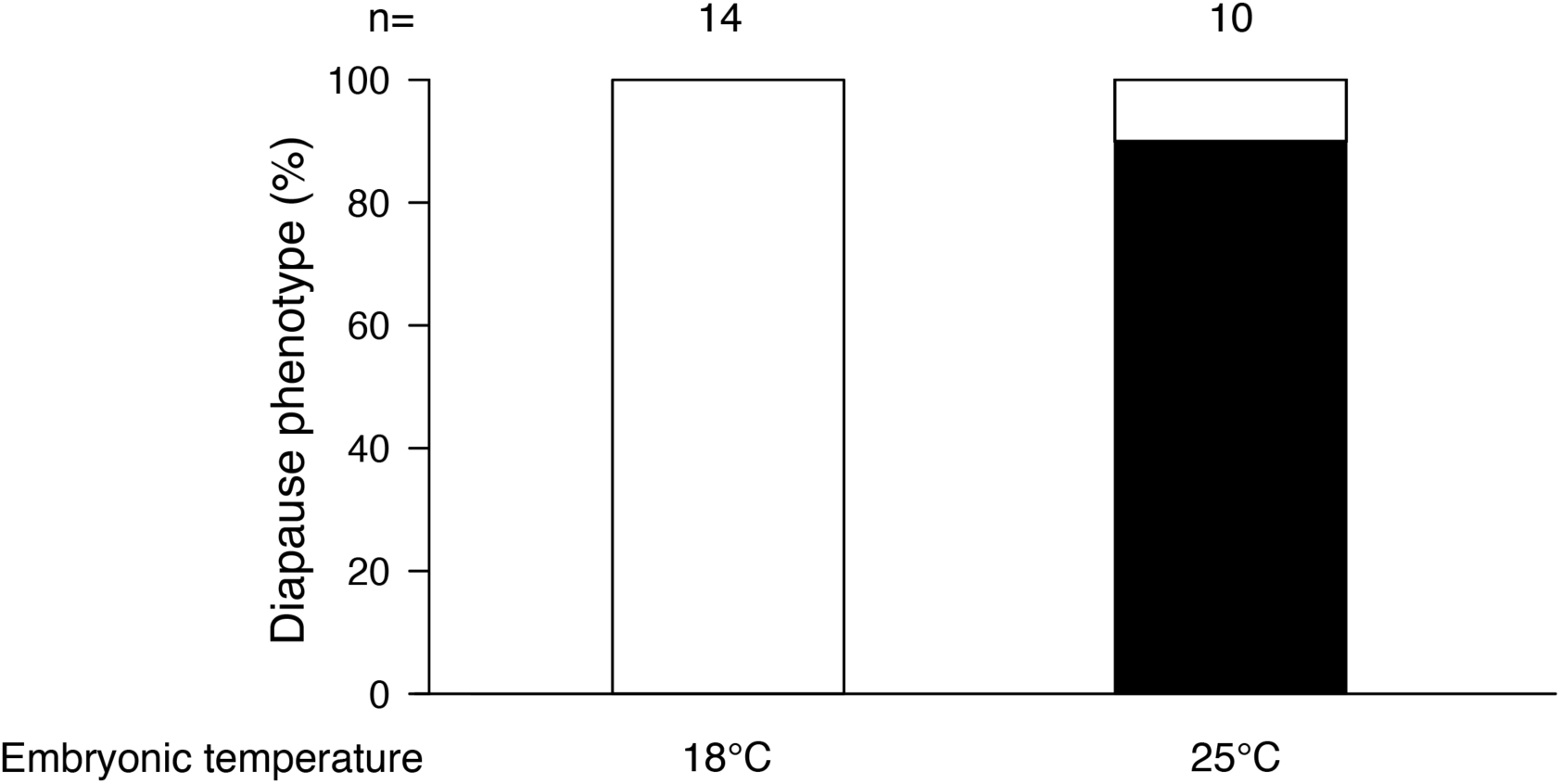
Diapause phenotype of the wild-type strain reared under continuous darkness. Eggs were incubated at 18℃ or 25℃ under continuous darkness until hatching. Hatched larvae were reared under continuous darkness as well. Black- and white-shaded parts show the proportions of females ovipositing diapause and non-diapause eggs, respectively.

**Figure 8.**
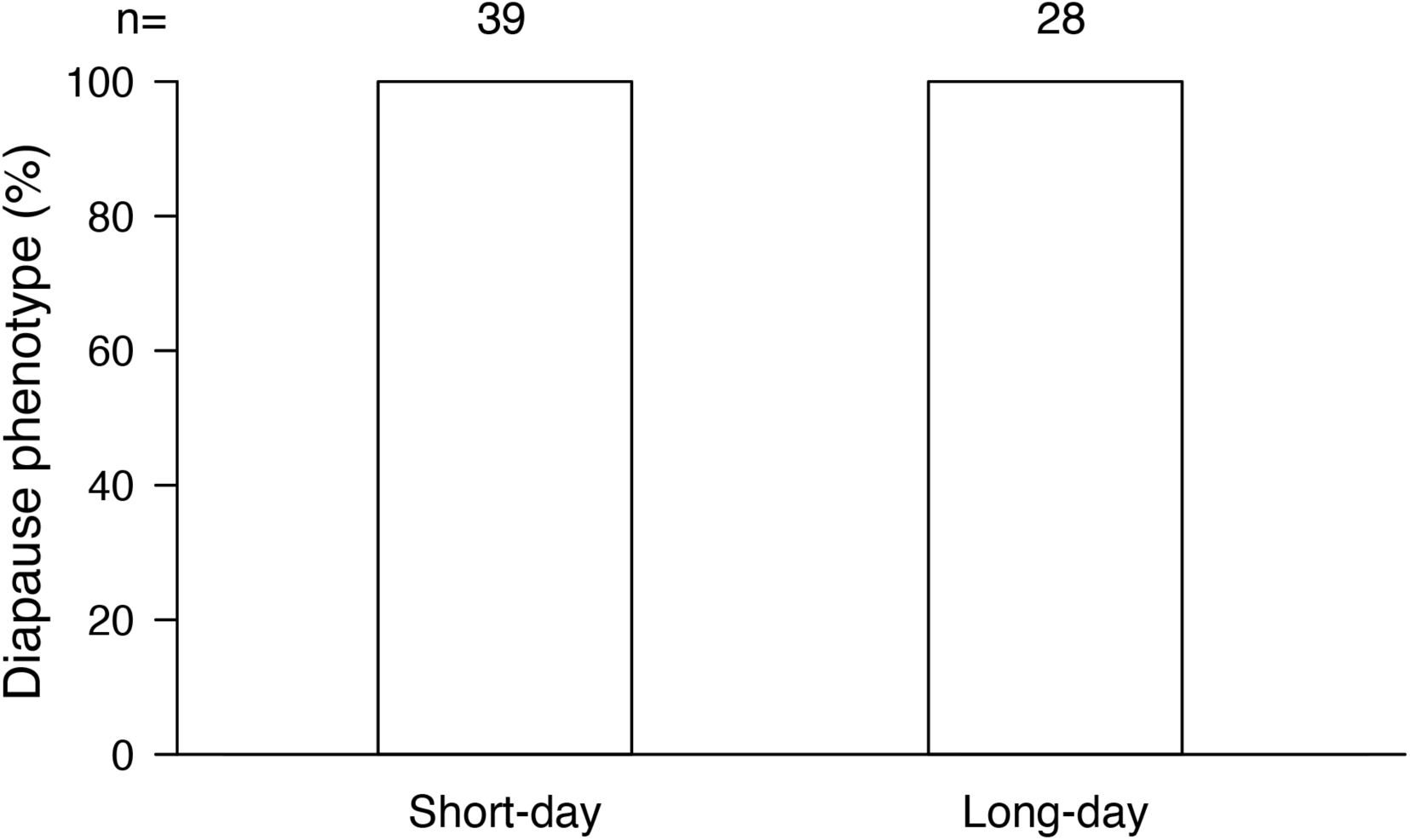
Diapause phenotype of the *cry1*/*tim* double-knockout strain during photoperiodic diapause induction in the larval stage. Eggs were incubated under continuous darkness at 25 ℃, and hatched larvae were reared under short-day (8L16D) or long-day (20L4D) conditions. All females oviposited non-diapause eggs under both photoperiodic conditions.

## 4. Discussion

### 4.1. Bombyx mori cry genes and the molecular clockwork

The generation of circadian rhythm in molecular clockworks differs among insects due to the presence or absence of the two *cry* genes. The present study revealed that the silkworm, *Bombyx mori*, also possesses two *cry* genes, *cry1* and *cry2*, with phylogenetic and structural analyses suggesting that these genes are *Drosophila*-type and mammalian-type, respectively. In addition to the clock genes *per*, *tim*, *Clk* and *cycle*, the presence of *cry-d* (*cry1*) and *cry-m* (*cry2*) implies that *B. mori* has a molecular clockwork similar to that of *D. plexippus*.

### 4.2. Expression profile of the B. mori cry1 gene

In the present study, the *cry1* gene exhibited rhythmic expression in the larval head of *B. mori*, peaking at early photophase. This expression patters aligns with findings in the adult brain and antennae of the African cotton leafworm, *Spodoptera littoralis* (Merlin et al., 2007), and in the larval and adult head of the cotton bollworm, *Helicoverpa armigera* (Yan et al., 2013; Liu et al., 2023). Our results are also consistent with a previous study involving *B. mori* embryos (Qiu et al., 2023a). Knockouts of core clock genes disrupted the temporal expression pattern of *cry1*, indicating that *cry1* expression is affected by the transcriptional feedback loop. The mRNA level of *cry1* increased in *Clk* and *cyc* knockout strains. In *D. melanogaster*, expression of *Clk* and *cry* (*cry-d*) is regulated by a second feedback loop composed of PDP1 and VRI proteins, which activate and suppress this expression, respectively, through VP-box (Cyran et al., 2003; Glossop et al., 2003). The upregulation of *cry1* expression in the *Clk* and *cyc* knockout strains in *B. mori* suggests that *cry1* expression is regulated by this second feedback loop, akin to the regulation of *cry* in *D. melanogaster*.

### 4.3. Bombyx mori cry1 is involved in the eclosion rhythm

The timing of hatching and adult eclosion in lepidopterans is under circadian clock control, as verified by clock gene knockout experiments in some species (Merlin et al., 2013; Brady et al., 2021; Chen et al., 2023; Liu et al., 2023; Wang et al., 2023). Previous studies in *B. mori* reported the loss of these rhythms in *per* and *tim* knockout strains (Ikeda et al., 2019; Nartey et al., 2021). Qiu et al. (2023a) reported the involvement of *cry1* in hatching rhythm through knockout experiments. The present study revealed that the *cry1* knockout strain lost its eclosion rhythm. Hence, the previous and current findings indicate that *cry1* is involved in both hatching and eclosion rhythms, similar to *per* and *tim*. Therefore, we conclude that the *B. mori cry1* gene is involved in circadian rhythm at the behavioral level. Several studies have reported the involvement of *cry1* in circadian rhythm in other lepidopterans. In the diamondback moth, *Plutella xylostella*, a *cry1* knockout strain exhibited arrhythmic locomotor activity (Chen et al., 2023). In *D. plexippus*, loss of functional *cry1* damped adult eclosion rhythm (Iiams et al., 2024). Therefore, *cry1* gene involvement in behavioral rhythms may be a common feature among lepidopterans.

### 4.4. Knockout of cry1 disrupts photoperiodism in B.mori

There are few studies about the involvement of *cry1* gene in photoperiodism in insects. In general, it has been suggested that CRY1 has an important role in photoperiodic time measurement as a photoreceptor (Saunders, 2012). In the cricket *M*. *siamensis*, Ueda et al. (2018) reported that knockdown of *cry1* affected the photoperiodic response in nymphal development. However, in crickets, CRY1 functions as a negative element, not as a photoreceptor (Tokuoka et al., 2017). Therefore, the involvement of photosensitive CRY in insect photoperiodism had not been investigated.

In the *cry1* knockout strain, nearly all females that experienced 25℃ conditions during the embryonic stage oviposited diapause eggs (Fig. 5), whereas those that experienced 18℃ conditions produced non-diapause eggs (Fig. 6). This phenotype differs from those of other clock gene knockout strains, which produce non-diapause eggs regardless of photoperiodic conditions. Additionally, we investigated diapause phenotype of the wild-type strain reared under constant darkness. As a result, when embryos were maintained under DD at 25℃, most of the resultant females produced diapause eggs; however, when embryos were maintained at 18℃ under DD, all resultant females produced non-diapause eggs (Fig. 7). Interestingly, these outcomes are similar to the diapause phenotypes of the *cry1* knockout strain, which appear to depend on embryonic temperature independent of larval photoperiods (Fig. 5 & 6). Based on these findings, we conclude that *cry1* is involved in photoreception of *B. mori* photoperiodism.

Notably, *cry1* knockout females responded weakly to photoperiod. Although most of females that experienced 25℃ during the embryonic stage produced diapause eggs, 20% of females oviposited non-diapause eggs when reared under long-day conditions, whereas females produced only diapause eggs when reared under short-day conditions (Fig. 5). Thus, it is possible that other photoreceptors compensate for circadian entrainment in the absence of *cry1*. Opsins expressed in the brain have been identified in various insects, with some suggested to contribute to circadian photoentrainment (Shimizu et al., 2001; Spaethe and Briscoe, 2005; Lampel et al., 2005; Velarde et al., 2005; Collantes-Alegre et al., 2018). For instance, Rh7 opsin expressed in the brain of *D. melanogaster* reacts to short-wave light, functioning in photoentrainment with CRY protein (Ni et al., 2017). Additionally, Shimizu et al. (2001) reported the expression of the long-wave type opsin *Boceropsin* in the larval brain of *B. mori*. The weak photoperiodic diapause induction of the *cry1* knockout strain may be attributed to the photoentrainment function of other photoreceptors expressed in the brain, such as *Boceropsin*. Photoreception via the brain has been reported previously in *B. mori* (Shimizu and Hasegawa, 1988). Immunocytochemistry using anti-*Drosophila* CRY antibody detected CRY1-like protein in the adult brain of *B. mori*, suggesting that BmCRY1 expression in the brain, akin to that in *D. melanogaster* (Sehadová et al., 2004). Futhermore, females of the *cry1*/*tim* double-knockout strain produced non-diapause eggs regardless of larval photoperiod, resembling the phenotype of the *tim* knockout strain, indicating that CRY1 functions upstream of the core feedback loop. Although further physiological studies are necessary to achieve an understanding of CRY1’s molecular function in *B. mori*, it is plausible that it functions as a photoreceptor in the brain during photoperiodic diapause induction.

## 5. Conclusion

In the present study, we established a *cry1* knockout strain and demonstrated the involvement of *cry1* in both circadian rhythm and photoperiodism in *B. mori*. Although several studies have implicated the circadian clock in insect photoperiodism, our understanding of its molecular underpinnings remains limited. Building upon our previous study (Tobita and Kiuchi 2022), we showed that knockout of canonical clock genes (*per*, *tim Clk*, *cyc*, and *cry1*), which play pivotal roles in the feedback loop of the circadian clock, influences photoperiodic diapause induction. This indicates the importance of this feedback loop in *B. mori* photoperiodism. Moreover, our study represents the first evidence of the involvement of *cry-d* in insect photoperiodism as a circadian photoreceptor.

## Acknowledgments

We thank the Institute for Sustainable Agro-ecosystem Services, The University of Tokyo, for facilitating mulberry cultivation and the Biotron Facility at the University of Tokyo for rearing the silkworms. We also thank Dr. Susumu Katsuma for useful advice and Dr. Kanako Hirota for the insect illustration used in the graphical abstract. We are grateful to Wakako Saito and Natsuki Nakashima for technical assistance during silkworm maintenance. This study was supported by JSPS KAKENHI grant numbers JP17H05047, JP20H02997, and 24K01768 to TK and JST SPRING (JPMJSP2108) to HT.

## Author contributions

HT and TK designed the study. HT and TK generated the knockout strains. HT conducted most of the experiments. HT and TK analyzed the data and wrote the manuscript. TK supervised the study.

## Supplementary figure

**Figure S1.**
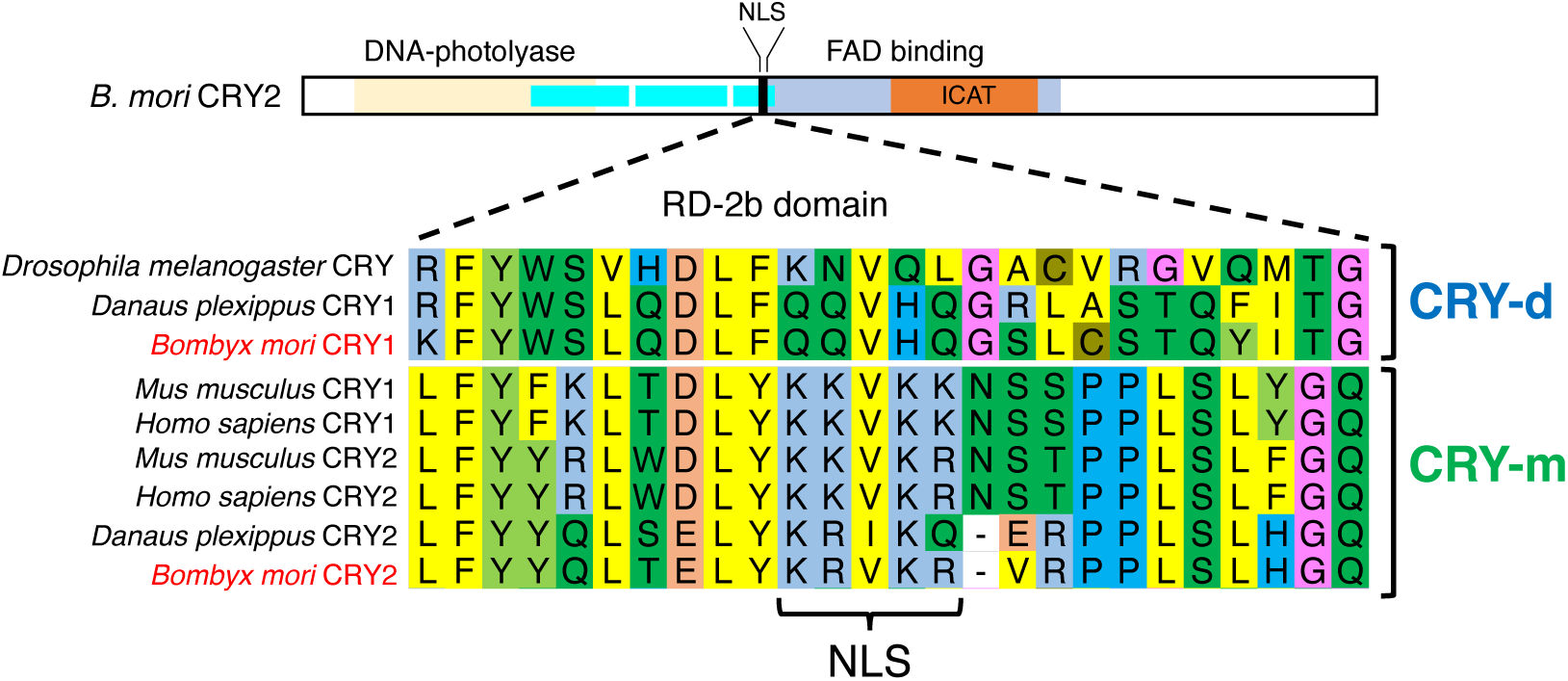
Alignment of CRY proteins around the nuclear localization signal (NLS). CRY amino acid sequences from *Drosophila melanogaster*, *Danaus plexippus*, *Bombyx mori*, *Mus musculus*, and *Homo sapiens* were aligned.

## Supplementary tables

**Table S1.** List of CRY protein homologs used in phylogenetic analysis.

| Species | Protein Name | Accession number |
| --- | --- | --- |
| *Drosophila melanogaster* | CRY | NP_732407.1 |
| *Anopheles gambiae* | CRY1 | ABB29886.1 |
| *Anopheles gambiae* | CRY2 | ABB29887.1 |
| *Gryllus bimaculatus* | CRY1 | BAX56238.1 |
| *Gryllus bimaculatus* | CRY2f | BAX56244.1 |
| *Modicogryllus siamensis* | CRY1 | BBD05665.1 |
| *Modicogryllus siamensis* | CRY2 | BBD05666.1 |
| *Myzus persicae* | CRY1 | AUN43313.1 |
| *Myzus persicae* | CRY2 | AUN43314.1 |
| *Antheraea pernyi* | CRY1 | AAK11644.1 |
| *Antheraea pernyi* | CRY2 | ABO38435.1 |
| *Bombyx mori* | CRY1 | NP_001182628.1 |
| *Bombyx mori* | CRY2 | NP_001182627.1 |
| *Danaus plexippus* | CRY1 | EHJ63675.1 |
| *Danaus plexippus* | CRY2 | EHJ74426.1 |
| *Apis mellifera* | CRY2 | NP_001077099.1 |
| *Tribolium castaneum* | CRY2 | NP_001076794.1 |
| *Xenopus laevis* | CRY1 | AAK94665.1 |
| *Xenopus laevis* | CRY2a | NP_001082139.1 |
| *Xenopus laevis* | CRY2b | AAK94667.1 |
| *Gallus gallus* | CRY1 | NP_989576.1 |
| *Gallus gallus* | CRY2 | NP_989575.1 |
| *Mus musculus* | CRY1 | NP_031797.1 |
| *Mus musculus* | CRY2 | NP_034093.1 |
| *Homo sapiens* | CRY1 | NP_004066.1 |
| *Homo sapiens* | CRY2 | NP_066940.3 |

**Table S2.**
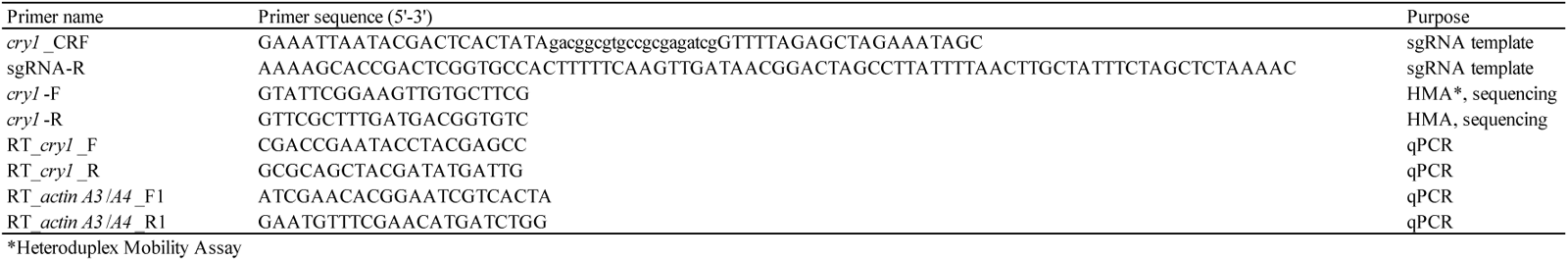
List of primers used in this study.

## Notes

### Competing Interest Statement

The authors have declared no competing interest.

## References

Allada, R., White, N.E., So, W.V., Hall, J.C., Rosbash, M., 1998. A mutant *Drosophila* homolog of mammalian clock disrupts circadian rhythms and transcription of *period* and *timeless*. Cell 93, 791–804. 10.1016/S0092-8674(00)81440-3

Ansai, S., Inohaya, K., Yoshiura, Y., Schartl, M., Uemura, N., Takahashi, R., Kinoshita, M., 2014. Design, evaluation, and screening methods for efficient targeted mutagenesis with transcription activator-like effector nucleases in medaka. Dev. Growth Differ. 56, 98–107. 10.1111/dgd.12104

Bassett, A.R., Tibbit, C., Ponting, C.P., Liu, J.L., 2014. Highly Efficient Targeted Mutagenesis of *Drosophila* with the CRISPR/Cas9 System. Cell Rep. 6, 1178–1179. 10.1016/j.celrep.2014.03.017

Beer, K., Helfrich-Förster, C., 2020. Model and Non-model Insects in Chronobiology. Front. Behav. Neurosci. 14, 1–23. 10.3389/fnbeh.2020.601676

Brady, D., Saviane, A., Cappellozza, S., Sandrelli, F., 2021. The Circadian Clock in Lepidoptera. Front. Physiol. 12. 10.3389/fphys.2021.776826

Ceriani, M.F., Darlington, T.K., Staknis, D., Más, P., Petti, A.A., Weitz, C.J., Kay, S.A., 1999. Light-dependent sequestration of TIMELESS by CRYPTOCHROME. Science. 285, 553–556. 10.1126/science.285.5427.553

Chen, S.P., Wang, D.F., Ma, W.F., Lin, X.L., Yang, G., 2023. Knockout of *cryptochrome 1* disturbs the locomotor circadian rhythm and development of *Plutella xylostella*. Insect Sci. 30, 1035–1045. 10.1111/1744-7917.13150

Collantes-Alegre, J.M., Mattenberger, F., Barberà, M., Martínez-Torres, D., 2018. Characterisation, analysis of expression and localisation of the opsin gene repertoire from the perspective of photoperiodism in the aphid *Acyrthosiphon pisum*. J. Insect Physiol. 104, 48–59. 10.1016/j.jinsphys.2017.11.009

Cortés, T., Ortiz-Rivas, B., Martínez-Torres, D., 2010. Identification and characterization of circadian clock genes in the pea aphid *Acyrthosiphon pisum*. Insect Mol. Biol. 19, 123–139. 10.1111/j.1365-2583.2009.00931.x

Cyran, S.A., Buchsbaum, A.M., Reddy, K.L., Lin, M.-C., Glossop, N.R.J., Hardin, P.E., Young, M.W., Storti, R. V, Blau, J., 2003. *vrille*, *Pdp1*, and *dClock* Form a Second Feedback Loop in the *Drosophila* Circadian Clock. Cell 112, 329–341.

Darlington, T.K., Wager-Smith, K., Ceriani, M.F., Staknis, D., Gekakis, N., Steeves, T.D.L., Weitz, C.J., Takahashi, J.S., Kay, S.A., 1998. Closing the circadian loop: CLOCK-induced transcription of its own inhibitors *per* and *tim*. Science. 280, 1599–1603. 10.1126/science.280.5369.1599

Egi, Y., Akitomo, S., Fujii, T., Banno, Y., Sakamoto, K., 2014. Silkworm strains that can be clearly destined towards either embryonic diapause or direct development by adjusting a single ambient parameter during the preceding generation. Entomol. Sci. 17, 396–399. 10.1111/ens.12073

Emery, P., So, W.V., Kaneko, M., Hall, J.C., Rosbash, M., 1998. CRY, a *Drosophila* clock and light-regulated cryptochrome, is a major contributor to circadian rhythm resetting and photosensitivity. Cell 95, 669–679. 10.1016/S0092-8674(00)81637-2

Emery, P., Stanewsky, R., Helfrich-Förster, C., Emery-Le, M., Hall, J.C., Rosbash, M., 2000. *Drosophila* CRY is a deep brain circadian photoreceptor. Neuron 26, 493–504. 10.1016/S0896-6273(00)81181-2

Glossop, N.R.J., Houl, J.H., Zheng, H., Ng, F.S., Dudek, S.M., Hardin, P.E., 2003. VRILLE feeds back to control circadian transcription of *Clock* in the *Drosophila* circadian oscillator. Neuron 37, 249–261. 10.1016/S0896-6273(03)00002-3

Goto, S.G., Nagata, M., 2022. The circadian clock gene (*Clock*) regulates photoperiodic time measurement and its downstream process determining maternal induction of embryonic diapause in a cricket. Eur. J. Entomol. 119, 12–22. 10.14411/EJE.2022.002

Hirayama, J., Sassone-Corsi, P., 2005. Structural and functional features of transcription factors controlling the circadian clock. Curr. Opin. Genet. Dev. 15, 548–556. 10.1016/j.gde.2005.07.003

Homma, S., Murata, A., Ikegami, M., Kobayashi, M., Yamazaki, M., Ikeda, K., Daimon, T., Numata, H., Mizoguchi, A., Shiomi, K., 2022. Circadian Clock Genes Regulate Temperature-Dependent Diapause Induction in Silkworm *Bombyx mori*. Front. Physiol. 13, 1–9. 10.3389/fphys.2022.863380

Iiams, S.E., Lugena, A.B., Zhang, Y., Hayden, A.N., Merlin, C., 2019. Photoperiodic and clock regulation of the vitamin A pathway in the brain mediates seasonal responsiveness in the monarch butterfly. Proc. Natl. Acad. Sci. U. S. A. 116, 25214–25221. 10.1073/pnas.1913915116

Iiams, S.E., Wan, G., Zhang, J., Lugena, A.B., Zhang, Y., Hardin, A.N., Merlin, C., 2024. Loss of functional *cryptochrome 1* reduces robustness of 24-hour behavioral rhythms in monarch butterflies. iSCIENCE 27. 10.1016/j.isci.2024.108980

Ikeda, K., Daimon, T., Sezutsu, H., Udaka, H., Numata, H., 2019. Involvement of the Clock Gene *period* in the Circadian Rhythm of the Silkmoth *Bombyx mori*. J. Biol. Rhythms 34, 283–292. 10.1177/0748730419841185

Ikeda, K., Daimon, T., Shiomi, K., Udaka, H., Numata, H., 2021. Involvement of the Clock Gene *Period* in the Photoperiodism of the Silkmoth *Bombyx mori*. Zoolog. Sci. 38, 523–530. 10.2108/zs210081

Ikeno, T., Ishikawa, K., Numata, H., Goto, S.G., 2013. Circadian clock gene *Clock* is involved in the photoperiodic response of the bean bug *Riptortus pedestris*. Physiol. Entomol. 38, 157–162. 10.1111/phen.12013

Ikeno, T., Numata, H., Goto, S.G., 2011. Photoperiodic response requires *mammalian-type cryptochrome* in the bean bug *Riptortus pedestris*. Biochem. Biophys. Res. Commun. 410, 394–397. 10.1016/j.bbrc.2011.05.142

Ikeno, T., Tanaka, S.I., Numata, H., Goto, S.G., 2010. Photoperiodic diapause under the control of circadian clock genes in an insect. BMC Biol. 8. 10.1186/1741-7007-8-116

Ingram, K.K., Kutowoi, A., Wurm, Y., Shoemaker, D.W., Meier, R., Bloch, G., 2012. The Molecular Clockwork of the Fire Ant *Solenopsis invicta*. PLoS One 7, 15–19. 10.1371/journal.pone.0045715

Kawamoto, M., Jouraku, A., Toyoda, A., Yokoi, K., Minakuchi, Y., Katsuma, S., Fujiyama, A., Kiuchi, T., Yamamoto, K., Shimada, T., 2019. High-quality genome assembly of the silkworm, *Bombyx mori*. Insect Biochem. Mol. Biol. 107, 53–62. 10.1016/j.ibmb.2019.02.002

Kogure, M., 1933. The influence of light and temperature on certain characters of the silkworm, Bombyx mori. J. Dep. Agric. Kyushu Imp. Univ. 4, 1–93. 10.5109/22568

Kotwica-Rolinska, J., Damulewicz, M., Chodakova, L., Kristofova, L., Dolezel, D., 2022. Pigment Dispersing Factor Is a Circadian Clock Output and Regulates Photoperiodic Response in the Linden Bug, *Pyrrhocoris apterus*. Front. Physiol. 13, 1–18. 10.3389/fphys.2022.884909

Kotwica-Rolinska, J., Pivarciova, L., Vaneckova, H., Dolezel, D., 2017. The role of circadian clock genes in the photoperiodic timer of the linden bug *Pyrrhocoris apterus* during the nymphal stage. Physiol. Entomol. 42, 266–273. 10.1111/phen.12197

Kumar, S., Stecher, G., Li, M., Knyaz, C., Tamura, K., 2018. MEGA X: Molecular evolutionary genetics analysis across computing platforms. Mol. Biol. Evol. 35, 1547–1549. 10.1093/molbev/msy096

Lampel, J., Briscoe, A.D., Wasserthal, L.T., 2005. Expression of UV-, blue-, long-wavelength-sensitive opsins and melatonin in extraretinal photoreceptors of the optic lobes of hawkmoths. Cell Tissue Res. 321, 443–458. 10.1007/s00441-004-1069-1

Liu, Xiaoming, Cai, L., Zhu, L., Tian, Z., Shen, Z., Cheng, J., Zhang, S., Li, Z., Liu, Xiaoxia, 2023. Mutation of the clock gene *timeless* disturbs diapause induction and adult emergence rhythm in *Helicoverpa armigera*. Pest Manag. Sci. 79, 1876–1884. 10.1002/ps.7363

Merlin, C., Beaver, L.E., Taylor, O.R., Wolfe, S.A., Reppert, S.M., 2013. Efficient targeted mutagenesis in the monarch butterfly using zinc-finger nucleases. Genome Res. 23, 159–168. 10.1101/gr.145599.112

Merlin, C., Lucas, P., Rochat, D., François, M.C., Maïbèche-Coisne, M., Jacquin-Joly, E., 2007. An antennal circadian clock and circadian rhythms in peripheral pheromone reception in the moth *Spodoptera littoralis*. J. Biol. Rhythms 22, 502–514. 10.1177/0748730407307737

Meuti, M.E., Stone, M., Ikeno, T., Denlinger, D.L., 2015. Functional circadian clock genes are essential for the overwintering diapause of the Northern house mosquito, *Culex pipiens*. J. Exp. Biol. 218, 412–422. 10.1242/jeb.113233

Mukai, A., Goto, S.G., 2016. The clock gene period is essential for the photoperiodic response in the jewel wasp *Nasonia vitripennis* (Hymenoptera: Pteromalidae). Appl. Entomol. Zool. 51, 185–194. 10.1007/s13355-015-0384-1

Naito, Y., Hino, K., Bono, H., Ui-Tei, K., 2015. CRISPRdirect: software for designing CRISPR/Cas guide RNA with reduced off-target sites. Bioinformatics 31, 1120–1123. 10.1093/bioinformatics/btu743

Nartey, M.A., Sun, X., Qin, S., Hou, C.X., Li, M.W., 2021. CRISPR/Cas9-based knockout reveals that the clock gene *timeless* is indispensable for regulating circadian behavioral rhythms in *Bombyx mori*. Insect Sci. 28, 1414–1425. 10.1111/1744-7917.12864

Ni, J.D., Baik, L.S., Holmes, T.C., Montell, C., 2017. A rhodopsin in the brain functions in circadian photoentrainment in *Drosophila*. Nature 545, 340–344. 10.1038/nature22325

Ota, S., Hisano, Y., Muraki, M., Hoshijima, K., Dahlem, T.J., Grunwald, D.J., Okada, Y., Kawahara, A., 2013. Efficient identification of TALEN-mediated genome modifications using heteroduplex mobility assays. Genes to Cells 18, 450–458. 10.1111/gtc.12050

Qiu, J., Cui, W., Zhang, Q., Dai, T., Liu, K., Li, J., Wang, Y., Sima, Y., Xu, S., 2022. Temporal transcriptome reveals that circadian clock is involved in the dynamic regulation of immune response to bacterial infection in *Bombyx mori* . Insect Sci. 1–16. 10.1111/1744-7917.13043

Qiu, J., Dai, T., Luo, C., Cui, W., Liu, K., Li, J., Sima, Y., Xu, S., 2023a. Circadian clock regulates developmental time through ecdysone and juvenile hormones in *Bombyx mori*. Insect Mol. Biol. 1–11. 10.1111/imb.12835

Qiu, J., Dai, T., Tao, H., Li, X., Luo, C., Sima, Y., Xu, S., 2023b. Inhibition of Expression of the Circadian Clock Gene *Cryptochrome 1* Causes Abnormal Glucometabolic and Cell Growth in *Bombyx mori* Cells. Int. J. Mol. Sci. 24, 5435. 10.3390/ijms24065435

Rubin, E.B., Shemesh, Y., Cohen, M., Elgavish, S., Robertson, H.M., Bloch, G., 2006. Molecular and phylogenetic analyses reveal mammalian-like clockwork in the honey bee (*Apis mellifera*) and shed new light on the molecular evolution of the circadian clock. Genome Res. 16, 1352–1365. 10.1101/gr.5094806

Rutila, J.E., Suri, V., Le, M., So, W.V., Rosbash, M., Hall, J.C., 1998. CYCLE is a second bHLH-PAS clock protein essential for circadian rhythmicity and transcription of *Drosophila period* and *timeless*. Cell 93, 805–814. 10.1016/S0092-8674(00)81441-5

Sato, Y., Shiomi, K., Saito, H., Imai, K., Yamashita, O., 1998. Phe-X-Pro-Arg-Leu-NH_2_ peptide producing cells in the central nervous system of the silkworm, *Bombyx mori*. J. Insect Physiol. 44, 333–342. 10.1016/S0022-1910(97)00140-6

Saunders, D.S., 2012. Insect photoperiodism: Seeing the light. Physiol. Entomol. 37, 207–218. 10.1111/j.1365-3032.2012.00837.x

Sehadová, H., Markova, E.P., Sehnal, F., Takeda, M., 2004. Distribution of circadian clock-related proteins in the cephalic nervous system of the silkworm, *Bombyx mori*. J. Biol. Rhythms 19, 466–482. 10.1177/0748730404269153

Shimizu, I., Hasegawa, K., 1988. Photoperiodic induction of diapause in the silkworm, *Bombyx mori*: location of the photoreceptor using a chemiluminescent paint. Physiol. Entomol. 13, 81–88. 10.1111/j.1365-3032.1988.tb00911.x

Shimizu, I., Matsui, T., Hasegawa, K., 1989. Possible Involvement of GABAergic Neurons in Regulation of Diapause Hormone Secretion in the Silkworm, *Bombyx mori*. Zoolog. Sci. 6, 809–812.

Shimizu, I., Yamakawa, Y., Shimazaki, Y., Iwasa, T., 2001. Molecular cloning of *Bombyx* cerebral opsin (boceropsin) and cellular localization of its expression in the silkworm brain. Biochem. Biophys. Res. Commun. 287, 27–34. 10.1006/bbrc.2001.5540

Spaethe, J., Briscoe, A.D., 2005. Molecular characterization and expression of the UV opsin in bumblebees: Three ommatidial subtypes in the retina and a new photoreceptor organ in the lamina. J. Exp. Biol. 208, 2347–2361. 10.1242/jeb.01634

Stanewsky, R., Kaneko, M., Emery, P., Beretta, B., Wager-Smith, K., Kay, S.A., Rosbash, M., Hall, J.C., 1998. The *cry^b^* mutation identifies cryptochrome as a circadian photoreceptor in *Drosophila*. Cell 95, 681–692. 10.1016/S0092-8674(00)81638-4

Takeda, S., Ogura, N., 1976. Induction of egg diapause by implantation of corpora cardiaca and corpora allata in *Bombyx mori*. J. Insect Physiol. 22, 941–944. 10.1016/0022-1910(76)90075-5

Tamai, T., Shiga, S., Goto, S.G., 2019. Roles of the circadian clock and endocrine regulator in the photoperiodic response of the brown-winged green bug *Plautia stali*. Physiol. Entomol. 44, 43–52. 10.1111/phen.12274

Tobita, H., Kiuchi, T., 2022. Knockouts of positive and negative elements of the circadian clock disrupt photoperiodic diapause induction in the silkworm, *Bombyx mori*. Insect Biochem. Mol. Biol. 149. 10.1016/j.ibmb.2022.103842

Tokuoka, A., Itoh, T.Q., Hori, S., Uryu, O., Danbara, Y., Nose, M., Bando, T., Tanimura, T., Tomioka, K., 2017. *cryptochrome* genes form an oscillatory loop independent of the *per* / *tim* loop in the circadian clockwork of the cricket *Gryllus bimaculatus*. Zool. Lett. 3, 1–14. 10.1186/s40851-017-0066-7

Truett, G.E., Heeger, P., Mynatt, R.L., Truett, A.A., Walker, J.A., Warman, M.L., 2000. Preparation of PCR-quality mouse genomic dna with hot sodium hydroxide and tris (HotSHOT). Biotechniques 29, 52–54. 10.2144/00291bm09

Tsuchiya, R., Kaneshima, A., Kobayashi, M., Yamazaki, M., Takasu, Y., Sezutsu, H., Tanaka, Y., Mizoguchi, A., Shiomi, K., 2020. Maternal GABAergic and GnRH/corazonin pathway modulates egg diapause phenotype of the silkworm *Bombyx mori*. Proc. Natl. Acad. Sci. U. S. A. 118. 10.1073/pnas.2020028118

Ueda, H., Tamaki, S., Miki, T., Uryu, O., Kamae, Y., Nose, M., Shinohara, T., Tomioka, K., 2018. *cryptochrome* genes mediate photoperiodic responses in the cricket *Modicogryllus siamensis*. Physiol. Entomol. 43, 285–294. 10.1111/phen.12258

Velarde, R.A., Sauer, C.D., Walden, K.K.O., Fahrbach, S.E., Robertson, H.M., 2005. Pteropsin: A vertebrate-like non-visual opsin expressed in the honey bee brain. Insect Biochem. Mol. Biol. 35, 1367–1377. 10.1016/j.ibmb.2005.09.001

Wang, D., Chen, J., Yuan, Y., Yu, L., Yang, G., Chen, W., 2023. CRISPR/Cas9-mediated knockout of period reveals its function in the circadian rhythms of the diamondback moth *Plutella xylostella*. Insect Sci. 30, 637–649. 10.1111/1744-7917.13139

Winfree, A.T., 1970. Integrated view of resetting a circadian clock. J. Theor. Biol. 28, 327–374. 10.1016/0022-5193(70)90075-5

Yan, S., Ni, H., Li, H., Zhang, J., Liu, X., Zhang, Q., 2013. Molecular cloning, characterization, and mRNA expression of two *Cryptochrome* genes in *Helicoverpa armigera* (Lepidoptera: Noctuidae). J. Econ. Entomol. 106, 450–462. 10.1603/EC12290

Yuan, Q., Metterville, D., Briscoe, A.D., Reppert, S.M., 2007. Insect cryptochromes: Gene duplication and loss define diverse ways to construct insect circadian clocks. Mol. Biol. Evol. 24, 948–955. 10.1093/molbev/msm011

Yuan, X.N., Luo, C., Zhao, Q.F., Zhong, S.Y., Hang, Q., Dai, T.M., Pan, Z.H., Sima, Y.H., Qiu, J.F., Xu, S.Q., 2023. The clock gene Cryptochrome 1 is involved in the photoresponse of embryonic hatching behavior in *Bombyx mori*. Arch. Insect Biochem. Physiol. 114, 1–17. 10.1002/arch.22046

Zhu, H., Sauman, I., Yuan, Q., Casselman, A., Emery-Le, M., Emery, P., Reppert, S.M., 2008. Cryptochromes define a novel circadian clock mechanism in monarch butterflies that may underlie sun compass navigation. PLoS Biol. 6, 0138–0155. 10.1371/journal.pbio.0060004

Zhu, H., Yuan, Q., Briscoe, A.D., Froy, O., Casselman, A., Reppert, S.M., 2006. The two CRYs of the butterfly. Curr. Biol. 16, 730. 10.1016/j.cub.2006.03.026

